# Characterization of Porcine Antibodies from Sequence Repertoire and Structural Data

**DOI:** 10.64898/2026.08.18.745626

**Authors:** Yoichi Kurumida, Yutaka Saito

## Abstract

Antibodies exhibit species-specific sequence and structural features that influence their antigen-recognition properties. Although several studies have investigated porcine antibodies, their repertoire and structural characteristics remain less well characterized than those of several other mammalian species. In this study, we analyzed public porcine heavy-chain repertoire sequencing data together with available antibody structural data to identify characteristic features of porcine antibodies. We found several residues enriched in porcine antibody framework regions, particularly at the base of heavy-chain complementarity-determining region 3 (CDR-H3). In particular, Arg101 and Glu123 were closely positioned in available structures and may influence CDR-H3 conformation at its base, whereas Pro120 may help constrain local backbone conformation. We also observed non-canonical cysteine usage in both framework region 1 and CDR-H3, which may contribute to structural diversity in the porcine repertoire. Finally, we evaluated the humanization potential of a porcine antibody using a human antibody language model and found that human-likeness increased after model-guided substitutions, although the resulting sequences did not exceed the T20 score threshold. Overall, these results indicate that porcine antibodies possess distinct sequence and structural features that may influence CDR-H3 properties and should be considered in future antibody analysis and engineering.

## Introduction

Antibody repertoires are highly diverse and contribute to host defense by recognizing a wide variety of foreign antigens^1^. The antigen-binding domain of an antibody is referred to as the variable region, which consists of complementarity-determining region (CDR) loops that directly participate in antigen recognition and framework regions (FRs) that support these loops as structural scaffolds^2^. The diversity of the antibody variable-region repertoire is generated primarily through V(D)J recombination, and somatic hypermutation (SHM)^3–5^. In particular, heavy-chain CDR3 (CDR-H3), which is formed at the V-D-J junction, is highly diverse in sequence and therefore plays a central role in antigen recognition^6^.

Antibodies are found widely across jawed vertebrates^7^. Although their overall architecture is broadly conserved, antibodies exhibit species-specific characteristics compared with human antibodies^8,9^. For example, shark antibodies include a stable single-domain format known as VNAR^10^, which contains four CDR loops. Structural differences in antibodies are also observed even among species within the same order. In cattle, which belong to the order Cetartiodactyla, a subset of antibodies possesses ultralong CDR-H3 regions and forms a characteristic “knob” structure at the tip of the CDR-H3 loop^11^. Likewise, some antibodies from camelids, also members of Cetartiodactyla, lack light chains and consist only of heavy chains^12^; in particular, the variable domain of these antibodies, known as VHH or nanobody, has been widely applied in therapeutics as a stable single-domain antibody fragment.

Pigs, another member of Cetartiodactyla, are known to use only four D-genes and five J-genes, resulting in a relatively limited sequence space generated by V-D-J combination^13^. Porcine antibodies may therefore exhibit characteristic sequence and structural features that help compensate for this restricted combinatorial diversity. Although several studies have reported large-scale sequencing analyses of porcine antibodies^14–17^, detailed sequence and structural analyses remain limited.

Here, we analyzed large-scale public sequencing data from porcine VH repertoires to investigate how porcine antibodies differ from those of humans. We further examined available antibody structures to assess how residues characteristic of pigs may influence structural features, considered the biological implications of these observations, and evaluated the potential applicability of porcine antibodies for human use.

## Results

### Overview of the porcine antibody repertoire

We analyzed porcine antibody VH sequences using NGS data from the Sequence Read Archive. After preprocessing, sequences derived from two pigs were annotated using IMGT/HighV-QUEST. A total of 321,425 sequences were included in the porcine dataset. The amino acid distributions of the framework are shown in Figure 1 and Figure S1 as sequence logos. Almost all sequences had framework regions of 25, 17, 38, and 11 amino acids in FR-H1, FR-H2, FR-H3, and FR-H4, respectively. CDR-H1 was predominantly 8 amino acids in length, whereas CDR-H2 was most frequently 8 or 10 amino acids in length

**Figure 1.**
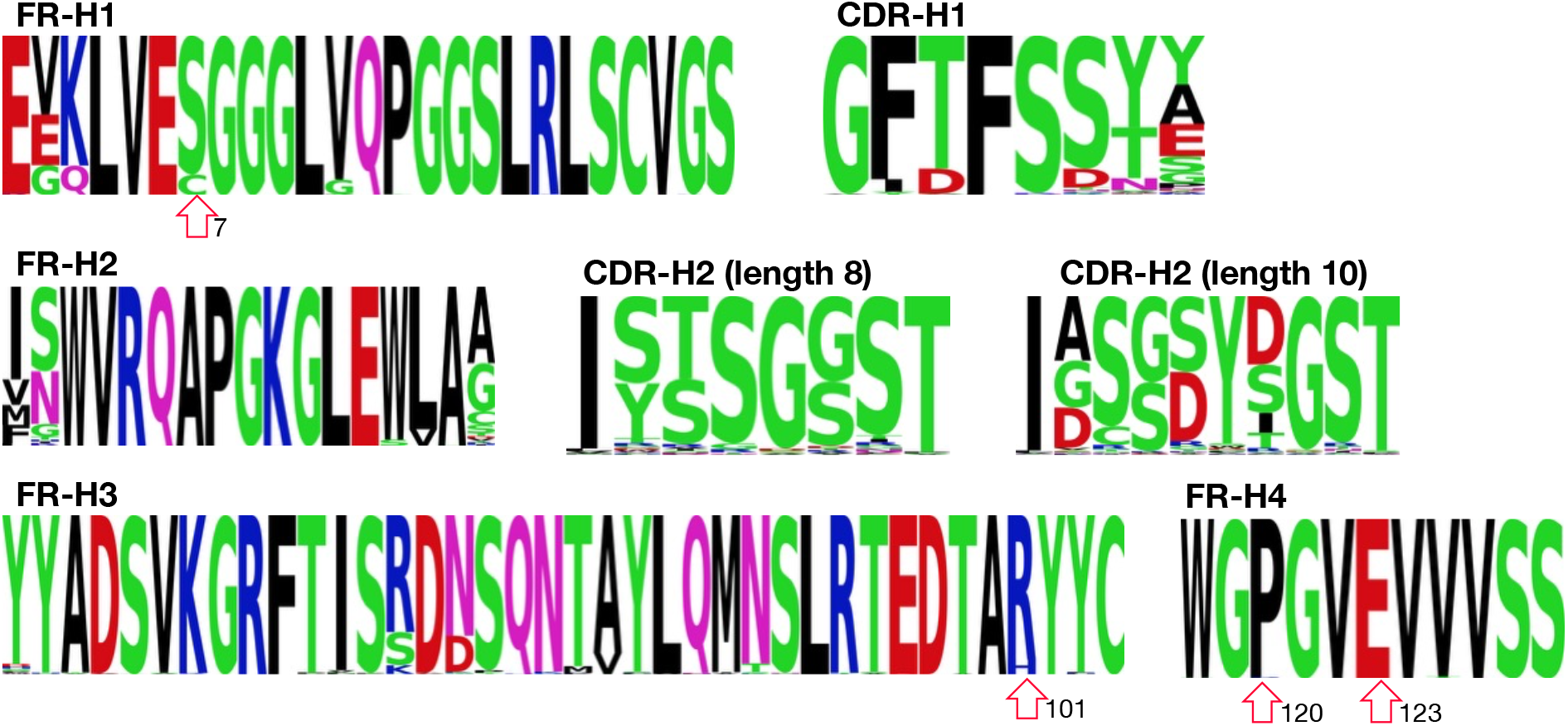
Distribution of porcine antibody sequences FR-H1, FR-H2, FR-H3, FR-H4, CDR-H1, and CDR-H2. Letters represent amino acid residues in the one-letter code, and heights indicate their frequencies at each position. Numbers indicate positions according to IMGT numbering

### FR-H residues characteristic of porcine antibodies cluster at the base of CDR-H3

Characteristic residues were identified in FR-H3 and FR-H4 at the base of CDR-H3. Specifically, these residues were Arg at IMGT position 101 in FR-H3 and Glu at position 123 in FR-H4^18^. In humans, these positions are more commonly occupied by Val and Leu, respectively (Figure S2). Examination of the three-dimensional structures showed that these two residues are located adjacent to each other between *β*-strands (Figure 2a,b). Analysis of the sequences of nine porcine antibodies deposited in the structural antibody database (SAbDab) showed that position 101 was occupied by Arg in eight antibodies and by His in one antibody. In our sequence dataset, a subset of sequences also had His at position 101. In contrast, all nine antibodies had Glu at position 123. Because these residues are located at the base of CDR-H3, they may influence the dynamics of CDR-H3 by restricting its conformational flexibility.

**Figure 2.**
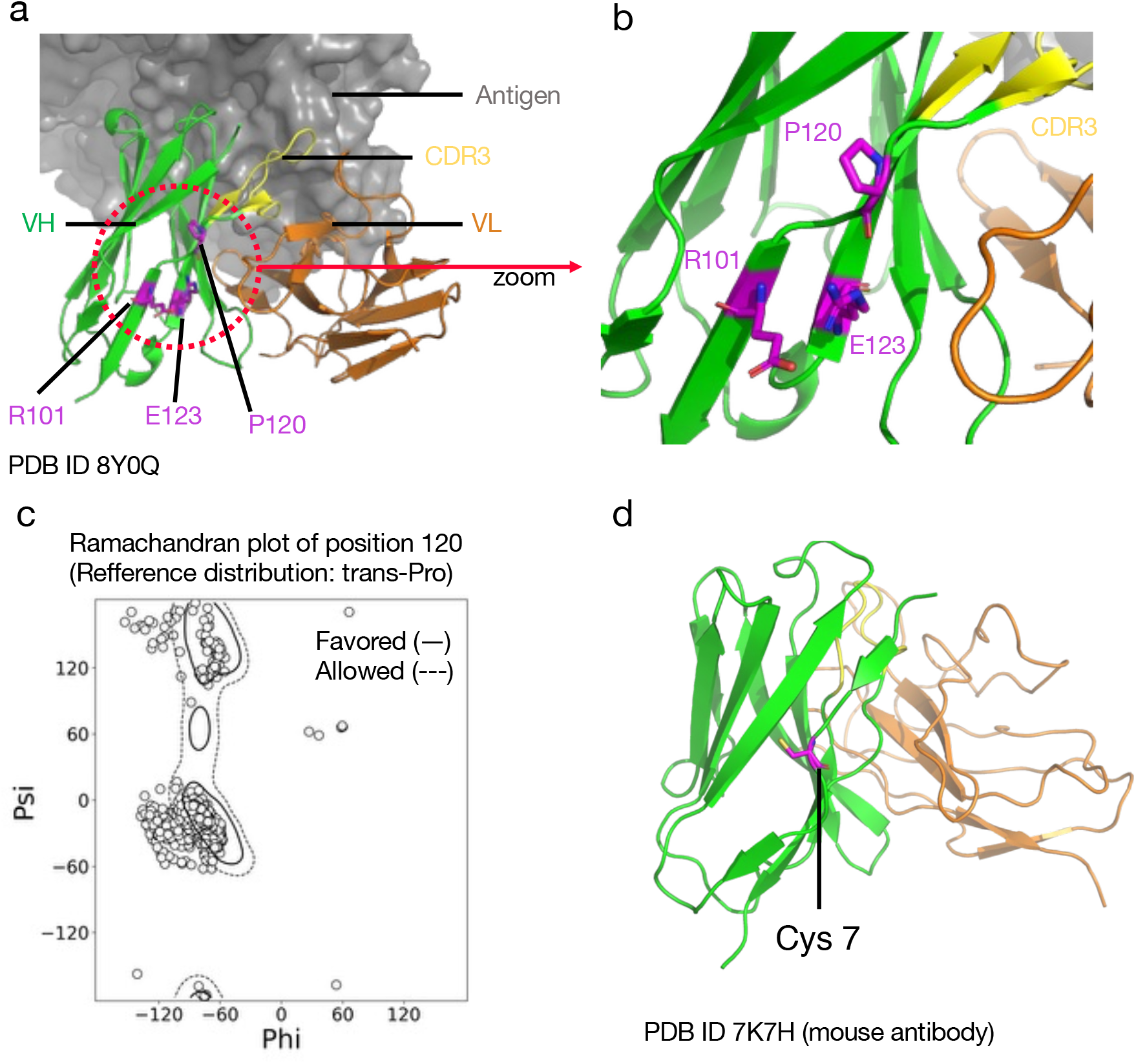
Characteristic residues of porcine antibodies around CDR-H3. a, Overview of the antibody-antigen complex structure. The VH domain is shown in green, the VL domain in orange, the antigen in gray, CDR-H3 in yellow, and the characteristic residues in magenta. b, Magnified view of panel a. c, Ramachandran plot for position 120. d, Structure around position 7.

In addition, Pro was observed at FR-H4 position 120, which is also located at the base of CDR-H3. In humans, this position is more frequently occupied by Gln (Figure S2). Because proline has a restricted range of backbone dihedral angles, it can impose conformational constraints on the surrounding structures deposited in SAbDab (Figure 2c). More than 70% of the observed conformations fell within the dihedral-angle range allowed for proline. This result suggests that substitution of the residue at position 120 with proline may constrain the local backbone dihedral angles and restrict motion to a stable conformational range in many antibodies, thereby contributing to conformational constraint at the base of CDR-H3.

### A subset of porcine antibodies contains a non-canonical cysteine in FR-H1

A subset of porcine antibodies contained a non-canonical free cysteine in FR-H1. Approximately 13% of antibodies had a cysteine at IMGT position 7, whereas serine was the dominant residue in both the human and porcine repertoires (Figure 1 and Figure S2). To examine the structural context of Cys7, we searched SAbDab for antibody structures carrying a cysteine at this position. Only one such structure was identified: a mouse-derived antibody (PDB ID: 7K7H; Figure 2d). In this structure, the cysteine at position 7 was located on the molecular surface rather than in the protein core, and it did not form a disulfide bond with another cysteine residue.

To explore the possible role of the free cysteine at position 7, we predicted post-translational modifications using the MusiteDeep web server^19^. The analysis predicted that the cysteine at position 7 could serve as a palmitoylation site. Palmitoylation is the covalent attachment of a palmitoyl group to a thiol group and is known to promote membrane localization by increasing hydrophobicity^20,21^. This raises the possibility that Cys at position 7 may influence the physicochemical properties of the antibody. At present, structural evidence supporting such a role remains limited.

### Porcine antibodies frequently contain non-canonical cysteines in CDR-H3

The features of CDR-H3 of the porcine antibodies, which contribute substantially to antigen binding, were analyzed. The mean CDR-H3 length was 15.7 amino acids, and the most frequent length was 17, followed by 18 and 15 (Figure 3a). A relatively high frequency was also observed for CDR-H3s of length 6, although separate analysis of the individual datasets showed that this increase was present in only one of the two datasets, suggesting that it may reflect individual variation or an immune response-specific bias (Figure S3). Analysis of the amino acids distributions of the most abundant CDR-H3 length groups, namely 15, 17, and 18, showed that the first and second residues at the N-terminal base of the loop were frequently Ala and Arg, respectively. At the C-terminal base of the loop, the third, second, and first residues from the end were frequently Met, Asp, and Leu, respectively. Within the loop region itself, amino acid residues commonly enriched in CDR-H3s, such as Gly, Tyr, Ser, and Ala, were also frequently observed in porcine antibodies (Figure 3b).

**Figure 3.**
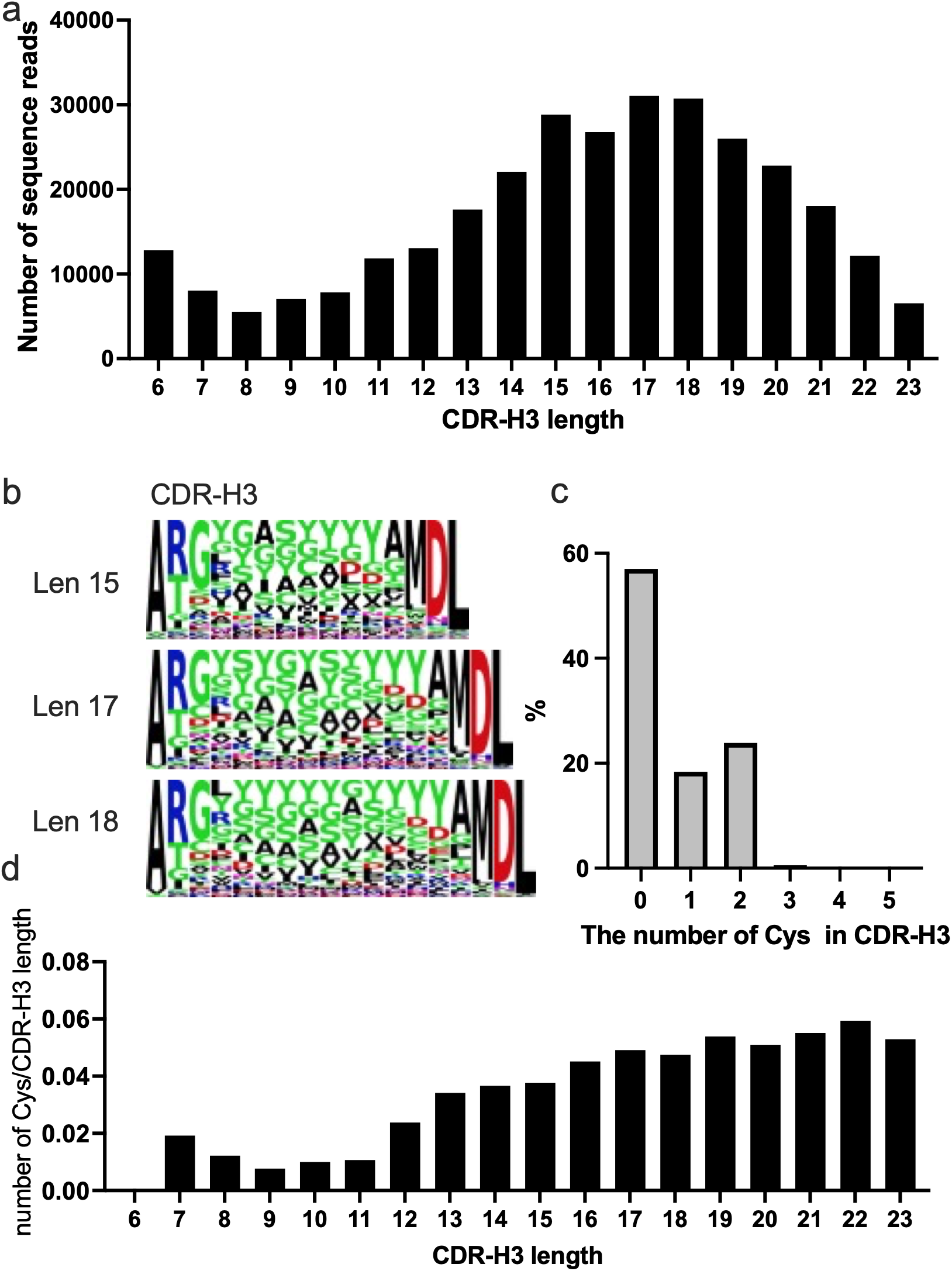
Analysis of the CDR-H3 region. a, Distribution of the CDR-H3 lengths. Only CDR-H3 lengths represented by 5,000 or more sequence reads are shown. b, Sequence logos of the three most frequent CDR-H3 lengths. c, Distribution of the cysteine residues in CDR-H3. d, Relationship between CDR-H3 length and the number of cysteine residues.

Next, we analyzed cysteine residues in CDR-H3 (Figure 3c). CDR-H3s containing no cysteine residues accounted for 57% of the repertoire, whereas 18% contained one cysteine and 24% contained two cysteines. Although less frequent, sequences containing three, four, or five cysteine residues were also observed. When the number of cysteine residues was analyzed as a function of CDR-H3 length, the number of cysteines increased with increasing CDR-H3 length (Figure 3d).

### Humanization of the porcine antibodies

To evaluate the potential applicability of porcine antibodies as human therapeutics, we performed humanization using a human antibody language model. A broadly neutralizing antibody VH against foot-and-mouth disease virus (FMDV), pOA2, was used for humanization^22^. Using the human antibody language model Sapiens^23^, we calculated the likelihood of each amino acid residue at every position (Figure 4a). Humanization was then performed by substituting each position with the residue showing the highest likelihood. Three humanization schemes were examined: FR-H only, FR-H with CDR-H1 and CDR-H2, and the entire variable region.

**Figure 4.**
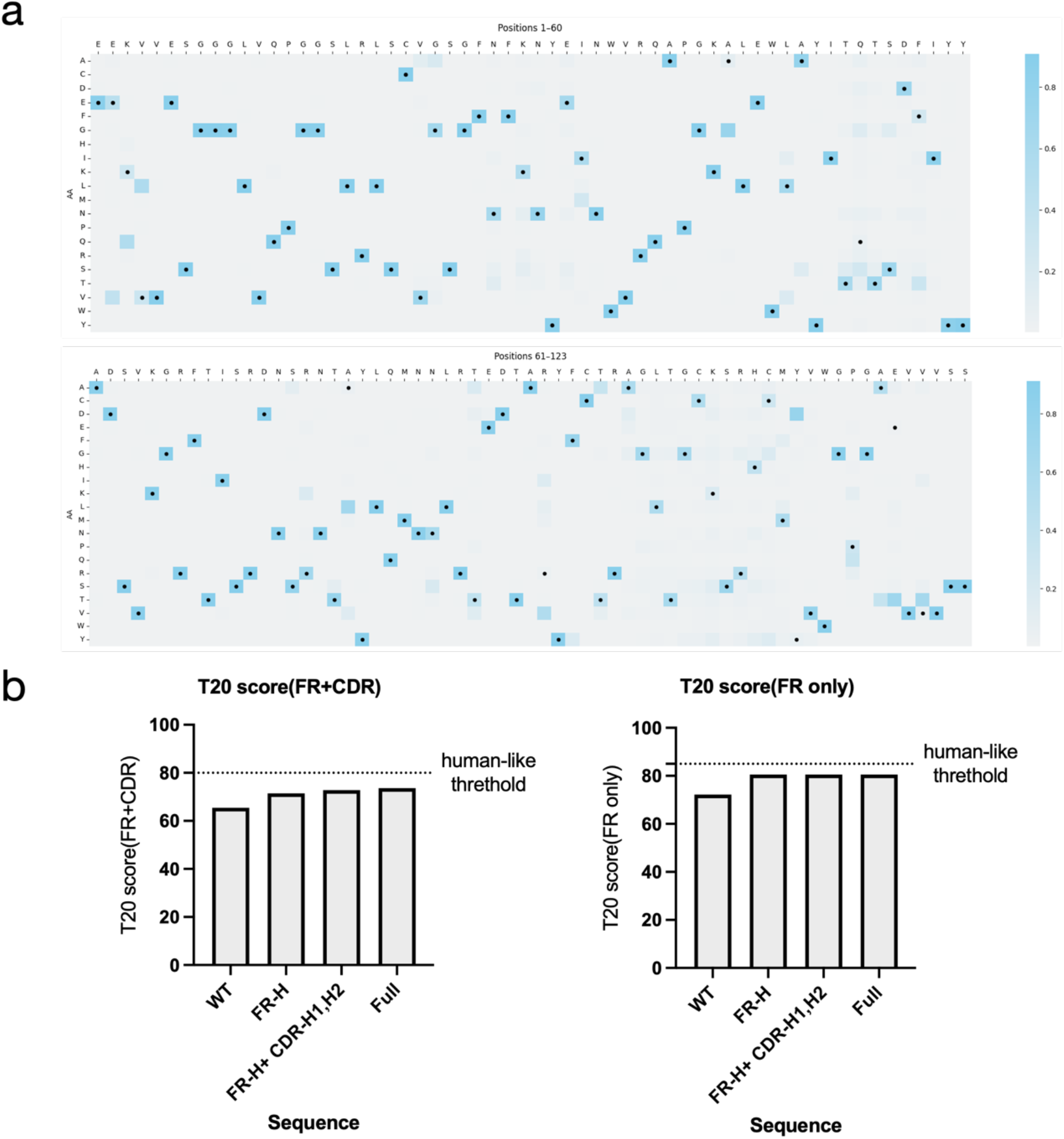
Evaluation of porcine antibodies using human antibody metrics. a, Score matrix calculated by the human antibody language model Sapiens. The black points indicate wild-type (WT) residues. b, Evaluation of human-likeness using T20 score.

As an evaluation of human-likeness, we calculated the T20 score using the web server (https://sam.curiaglobal.com/t20/)^24^, an alignment-based metric of similarity to human antibodies (Figure 4b). The T20 score is defined as the average sequence identity to the top 20 most similar human-derived antibody sequences. Overall human-likeness increased as the humanized region expanded in the Sapiens-based designs; however, none of the designs exceeded the threshold defined by the T20 score. This result suggests that porcine antibody features may complicate direct translation to human therapeutics.

## Discussions

In this study, we identified several residues characteristic of porcine antibodies by analyzing large-scale public sequence and structural datasets. The salt bridge formation observed in FR-H3 and FR-H4, as well as the substitution to proline in FR-H4, may contribute to conformational constraint at the base of CDR-H3 by restricting local dynamics. Although CDR-H3 plays a critical role in antigen recognition in antibodies^25^, it may be particularly important in porcine antibodies because the diversity of V, D, and J genes is relatively limited, suggesting that the diversity generated at the V(D)J recombination junctions accounts for a substantial proportion of the overall diversity in the variable region^26^. Therefore, CDR-H3 may play an especially important role in antigen recognition in pigs, and these features may also contribute to conformational constraint at the base of CDR-H3.

In addition, the presence of non-canonical cysteines in CDR-H3 may also contribute to conformational constraint by restricting loop flexibility through disulfide bond formation. In chicken antibodies, increased CDR-H3 length has been reported to be associated with an increased number of cysteines^28^, and a similar trend was observed in porcine antibodies in this study (Figure 3d). In human antibodies, only approximately 10% CDR-H3 contain one or two non-canonical cysteines^27^, with these cysteines occurring more frequently in longer CDR-H3 loops. Nearly 18% of porcine antibody sequences contained only a single cysteine within CDR-H3. Although free cysteines are generally thought to have the potential to impair folding efficiency, the biological significance of this observation remains unclear.

We also applied an existing human antibody language model to humanize porcine antibodies and evaluated the results using other alignment-based methods, which suggested a certain degree of humanization potential. Because pigs are also hosts of zoonotic pathogens^29^, the ability to humanize porcine antibodies could provide an efficient and relatively low-risk approach for therapeutic antibody development.

One limitation of this study is that the analysis was performed using only data from Landrace pigs born from the same sow, which may include genetic bias as well as bias at the breed level. Several studies have reported large-scale sequence analyses of porcine antibodies, including publicly available SRA datasets^15,17^. One of these studies analyzed antibodies at the single-cell level and therefore included paired analyses with VL, but the dataset size was limited^17^. Another study involved immunization with inactivated FMDV serotype Asia1 vaccine, which may have introduced bias toward particular sequences^17^. Although the dataset used in this study corresponded to the non-immunized control group in the original publication ^30^. The authors did not state that the pigs were raised under germ-free conditions; therefore, it remains possible that they had already acquired immune experience.

In conclusion, this study revealed several features characteristic of porcine antibodies. These findings suggested that porcine antibodies may influence CDR-H3 conformation, which appears to be a major source of antibody diversity in this species. We hope that the insights obtained in this study will contribute to engineering approaches for antibodies, which are important not only as research and diagnostic tools, but also as therapeutic agents.

## Methods

### Datasets

Two porcine antibody VH sequence datasets were retrieved from the NCBI Sequence Read Archive (SRA; SRR12072415 and SRR12072416). Briefly, the datasets were obtained from two pigs derived from the same Landrace sow. Peripheral blood mononuclear cells were collected at 3 weeks of age. After RNA extraction and reverse transcription, antibody sequences were amplified using Fv-specific primers. Sequencing was performed on the Illumina MiSeq platform using paired-end 2×250 bp reads. Duplicate sequences were not removed prior to analysis. Human antibody VH sequence retrieved from the abYsis web server (accessed on August 14, 2026)^31^.

### Sequence preprocessing

We processed the raw sequencing data to obtain high-quality antibody sequences. First, adaptor sequences and low-quality bases (Q < 20) were removed using the fastp software^32^. Paired-end reads were then merged using FLASH2^33^. Sequences containing ambiguous bases (“N”) were discarded. The filtered nucleotide sequences were translated into amino acid sequences and annotated using the IMGT/HighV-QUEST web server^34^. Only productive sequences were retained, and sequences containing stop codons or missing annotations(“nan”) were excluded.

### Analysis of Dihedral Angles

To evaluate antibody backbone conformations, we calculated dihedral angles using the Python API of PyMOL. Antibody structures were retrieved from SAbDab^35^, and only structures with a resolution of 1.5 Å or better were included. Structures in which the heavy and light chains were not separately annotated, or in which coordinates for the residue of interest were missing, were excluded. The resulting angles were visualized as a Ramachandran plot. As a reference for the natural distribution of dihedral angles, we used the cctbx package in Python^36^.

### Post-translational modification prediction

Potential post-translational modifications were predicted using the MusiteDeep web server^19^ (accessed on February 23, 2026). All available PTM-specific prediction models were applied to the VH amino acid sequence, and all other parameters were kept at their default settings. The query sequence is provided in Supplementary Data 1.

### Humanization

The protein language model Sapiens was used to humanize a porcine sequence^22,23^. For the sequence, positional scores were calculated with Sapiens using single-pass prediction without any mask tokens, and the amino acid with the highest score at each position was selected as the humanized residue.

### Visualization

Sequence logos were generated using the Logomaker package in Python. Other graphs were prepared using GraphPad Prism or the matplotlib and seaborn packages in Python

## Acknowledgements

This work was partly supported by JSPS KAKENHI (Y.K., 24K20894). The computations were partly performed on the NIG supercomputer at ROIS National Institute of Genetics, Miyabi supercomputer at the University of Tokyo, and the Wisteria supercomputer at the University of Tokyo through the HPCI System Research Project (hp240164, hp250106). We thank the developers of OpenAI’s ChatGPT for assistance with English editing of the paper.

## Author contributions

Y.K.: Conceptualization, Formal analysis, Methodology, Visualization, Writing—original draft, and Writing—review & editing. Y.S.: Methodology and Writing—review & editing

## Competing interests

The authors declare no competing interests.

## Notes

### Competing Interest Statement

The authors have declared no competing interest.

