## Supplementary figures and data for "Characterization of Porcine Antibodies from Sequence Repertoire and Structural Data"

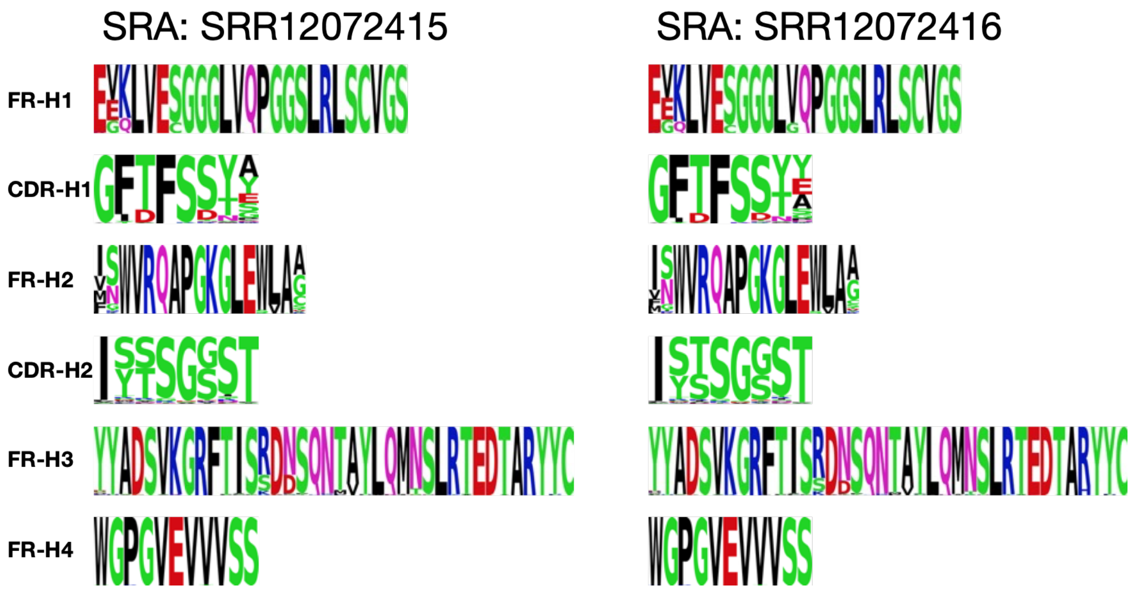


Supplementary Figure 1. Independent analysis of the porcine sequence data.

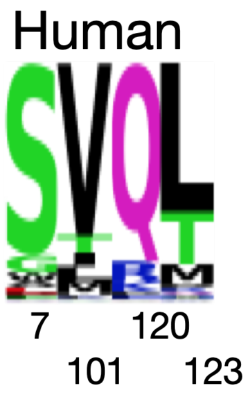


Supplementary Figure 2. Distribution of residues at selected positions in human antibody VH.

SRA: SRR12072415 SRA: SRR12072416

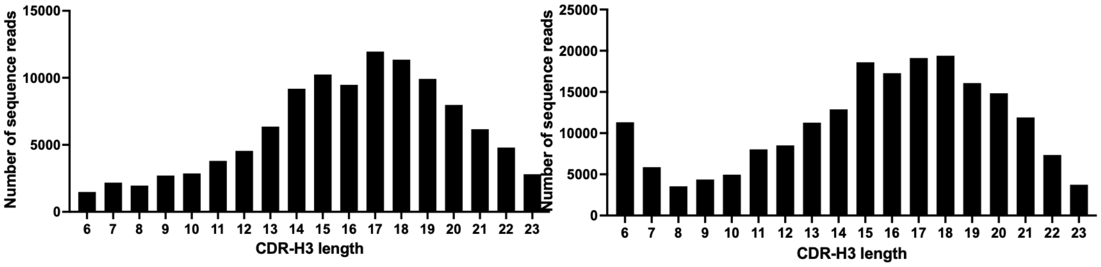


Supplementary Figure 3. Independent analysis of the porcine CDR-H3.

Supplementary Data 1. The query sequence used for MusiteDeep prediction.

EEKLVECGGGLVQPGGSLRLSCVGSGFTFSSYSMSWVRQAPGKGLEWLAIIYSSGSSTYYAD
SVKGRFTISRDNSQNTAYLQMNSLRTEDTARYYCATGLSSWGPGVEVVVSS
